# Decision benchmarking *reveals* when protein foundation models add experimental value

**DOI:** 10.64898/2026.10.13.758966

**Authors:** Wenze Ding, Chenlin Lu, Yao Li, Haipeng Gong, Zhiyong Wang

## Abstract

Protein engineering is shifting from model-based variant scoring to deciding what to measure next. This exposes a limitation of fixed-dataset benchmarks, which rank predictors without showing when their advantage survives as evidence accumulates. We therefore introduce decision benchmarking across 217 ProteinGym assays, 2.47 million variants and 95 predictors to evaluate whether models and measurements improve decisions at matched cost. Zero-shot accuracy explained less than one-fifth of the variation in discovery after calibration. Calibrated priors exceeded budget-matched additivity at 20–100 measurements, but the advantage narrowed as component measurements accumulated. Measured additivity then overtook zero-shot models after 50–100 measurements for top-variant discovery and around 500 for global ranking. The same principle showed that double-mutant measurements improved prediction in models using mutation identity, whereas prediction-guided acquisition improved discovery. Evidence-based routing improved strategy selection. Decision benchmarking thus identifies which strategy adds value given the information already available and the decision that follows.

## Main

Protein engineering searches vast sequence spaces for variants with improved activity, stability, binding and other experimentally measured properties. Directed evolution proceeds through mutation, measurement and selection^1–3^, often using only tens of measured variants to guide each round^4–7^. Deep mutational scanning^8,9^ and resources such as ProteinGym^10^ and FLIP^11^ have mapped sequence–function landscapes across diverse proteins^12–14^, enabling systematic evaluation of these experimental choices^15,16^. The practical question is how effectively a predictor directs a limited measurement budget towards the experimental objective. A useful strategy must make productive use of measurements already collected and identify which candidates are worth testing next.

Protein foundation models learn reusable representations from sequence^17–26^ and structural information^27–33^, complementing family-specific evolutionary information^34–37^ and encoding interpretable protein features^38^. Benchmarks such as ProteinGym^10^ and ProteinBench^39^ assess these models across proteins and tasks^40–43^. Zero-shot scores provide predictions before target-assay measurements; once measurements become available, scores can be calibrated, supervised models fitted and subsequent variants selected through experimental feedback^4,5,44–47^. Each step expands the strategies a campaign can choose between. Comparisons conducted with different candidate pools, budgets and objectives leave unclear when accumulated measurements change the preferred strategy. The unresolved quantity is the measurement depth at which empirical information overtakes a model prior, and how that depth changes with the experimental objective. Locating it requires comparing strategies as their available information grows under matched experimental conditions.

Measured biological controls make this question testable. These controls exploit known biological relationships, as measured component single-mutant effects can approximate a double-mutant phenotype through their additive expectation^48–51^. The information available to this control grows with component coverage, making measurement depth part of the comparison with a model prior. A predictor can recover these component effects and rank double mutants accurately without capturing epistasis, the departure of combined mutation effects from additivity^51,52^. Claims of interaction learning must therefore exceed an explicit additive control supplied with a stated amount of component information. The same requirement extends to acquisition signals. Uncertainty or disagreement may identify unusual variants, but their experimental value is established only if measuring those variants improves subsequent prediction or candidate selection at the same cost.

Here we establish decision benchmarking across 217 ProteinGym substitution assays, 2,465,767 variants and 95 predictors, comparing complete experimental strategies as their available information and measurement budgets change. We locate a measurement-depth crossover at which measured additivity overtakes zero-shot references, occurring earlier for top-variant discovery than for global ranking. At small matched campaign budgets, calibrated priors retain a substantial advantage over additivity. These comparisons show that predictor choice depends jointly on the information already available and the decision being optimized. We then test how additional measurements alter prediction and candidate selection, and integrate this evidence into assay-level starting strategies. In summary, these analyses connect model evaluation to experimental design by showing when strategy choice should change as experimental information accumulates.

## Results

### Decision benchmarking evaluates complete experimental strategies

We defined the evaluation unit as a complete strategy operating within one assay. Each strategy was specified by its candidate pool, measurement budget and available information. The candidate pool defined which variants could be considered for testing, while the measurement budget defined how many assay outcomes were available to guide the strategy. Available information included model priors supplied by precomputed zero-shot scores before any assay measurements were used. Within each comparison, strategies shared the same measured variants and held-out candidates, then ranked variants, calibrated scores, fitted supervised models or selected further measurements (Fig. 1a). This design follows the choices a campaign makes as information accumulates and evaluates how each strategy turns that information into an experimental action.

**Figure 1.**
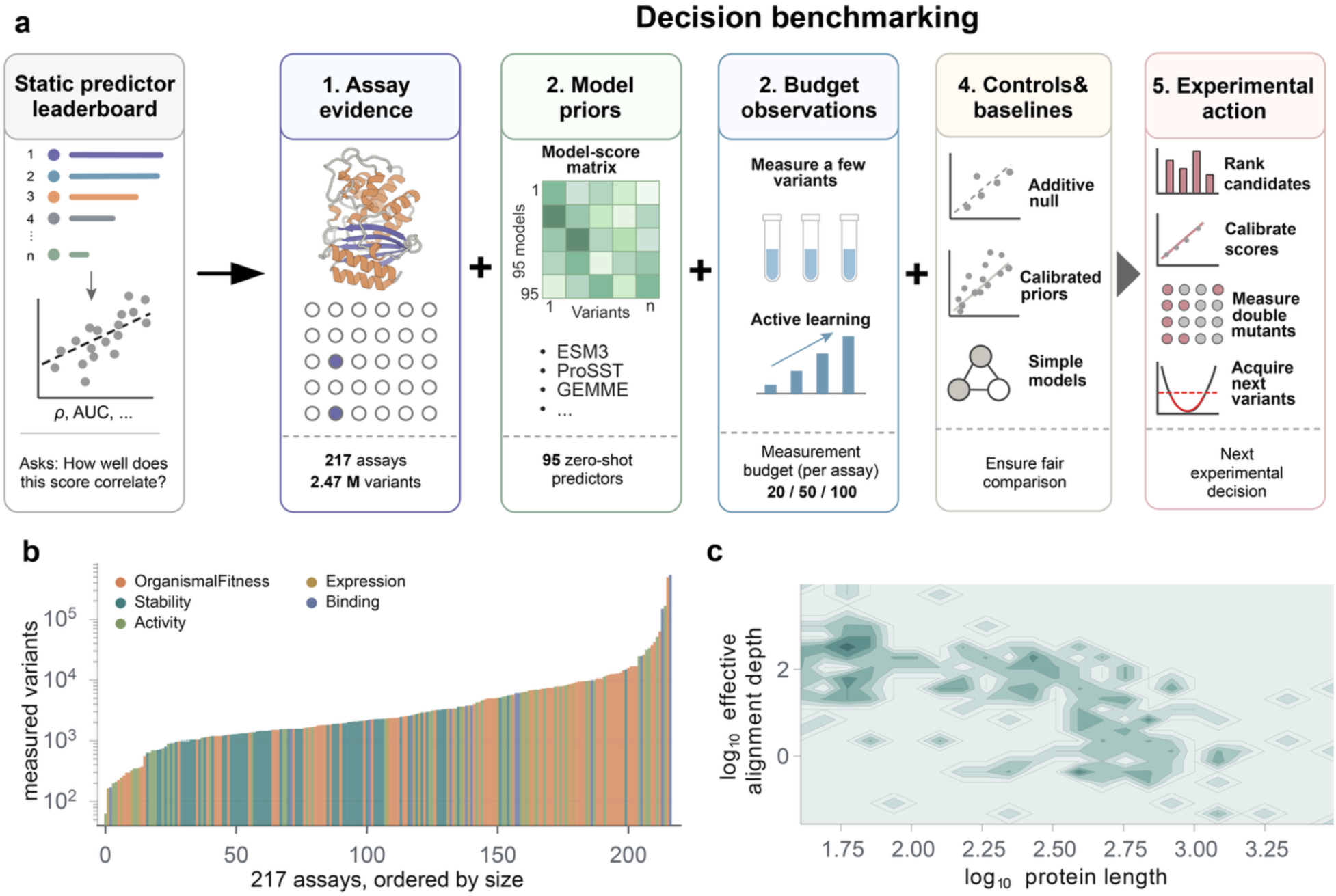
Decision benchmarking for data-efficient protein engineering. **a**, The evaluation unit combines an assay and candidate pool, model priors, budgeted observations, matched controls and an experimental action. The benchmark includes 217 assays, 2.47 million measured variants and 95 zero-shot predictors, with budgets of 20, 50 or 100 measurements per assay. Actions include ranking candidates, calibrating scores, measuring double mutants and acquiring subsequent batches. **b**, Variant counts for the 217 assays, ordered by size and coloured by phenotype class; the logarithmic axis spans four orders of magnitude. **c**, Two-dimensional density of effective alignment depth against protein length, both on base-10 logarithmic scales.

The benchmark comprises 217 substitution assays, 2,465,767 measured variants and 95 predictors spanning sequence, structural and evolutionary information (Supplementary Figs. 1 and 2). 72% of variants carry at least two substitutions, providing the multi-mutant coverage needed to compare model priors with measured component effects (Fig. 1b; Supplementary Tables 1 and 2). Protein lengths range from 37 to more than 3,000 residues and alignment depths span four orders of magnitude (Fig. 1c). All assays supported zero-shot and few-measurement evaluations, while 69 contained component-resolved double mutants, 11 contained higher-order variants, and 27 of 28 selected assays supported the common-pool active-learning comparison (Supplementary Fig. 1; Supplementary Tables 3 and 4).

Performance was summarized within assay before aggregation, giving each assay equal weight despite differences in library size. Deterministic splits were shared across strategies at each budget to preserve matched comparisons. To compare experimental strategies, we fixed the candidate pool, measurement budget and available information, then evaluated the decision made by each strategy.

### Measurements and objectives reshape the value of a model prior

We first evaluated the priors available before target-assay measurement on two objectives. Absolute Spearman correlation assessed global ordering, whereas top-variant discovery measured the recovery of high-performing variants in a short testing list. These objectives distinguish a useful landscape ranking from a useful experimental shortlist. VenusREM^53^, ProSST-2048^32^ and S3F-MSA^54^ led global ranking, with mean absolute Spearman correlations of 0.536, 0.523 and 0.516 across 217 assays (Fig. 2a,b; Supplementary Table 5). Within-family variation exceeded between-family differences for protein language models, and performance varied across assays (Fig. 2c). For discovery, ProSST-2048 achieved the highest top-20 enrichment of measured top-1% variants, 5.86-fold over random, followed by VenusREM at 5.62-fold and ESM3 at 5.55-fold (Fig. 2d; Supplementary Tables 6 and 7). The leading group persisted across robustness analyses (Supplementary Fig. 2). These scores provide useful starting priors before assay-specific measurements create opportunities to adapt them and build competing predictors.

**Figure 2.**
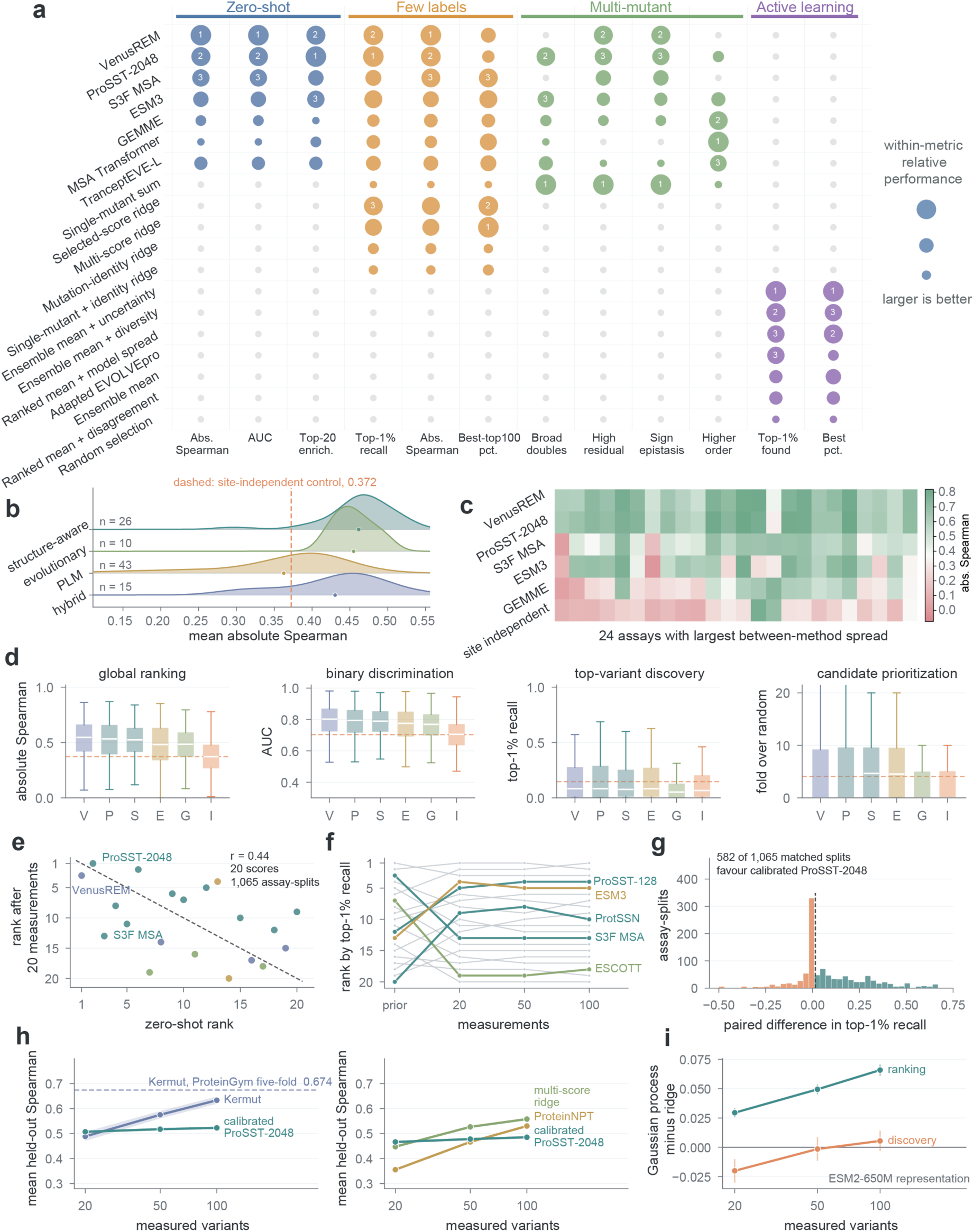
Measurement budget and objective determine the value of a model prior. **a**, Strategies compared across decision metrics. Circle area indicates relative performance within each metric; numbers identify the three highest-ranked strategies. Grey circles denote undefined comparisons. **b**, Distributions of mean absolute Spearman correlation across methods in each family; points mark family means, labels give method counts and the dashed line marks the site-independent control (0.372). **c**, Six representative scores across the 24 assays with the greatest between-method variation. **d**, Assay-level distributions of four zero-shot metrics. Boxes span the interquartile range, white lines mark medians and whiskers extend to the most extreme values within 1.5 times the interquartile range; outliers are omitted. Dashed lines mark the site-independent mean. V, VenusREM; P, ProSST-2048; S, S3F-MSA; E, ESM3; G, GEMME; I, site independent. **e**, Zero-shot and 20-measurement discovery ranks for 20 scores. The annotated Pearson r = 0.44 relates the underlying method-level mean zero-shot correlations and calibrated top-1% recall values; ranks are shown for comparison. **f**, Top-1% recall ranks before measurement and at budgets of 20, 50 and 100. **g**, Paired difference in recall between calibrated ProSST-2048 and measured additivity across 1,065 assay-splits; 582 favour calibration. **h**, Mean held-out Spearman correlation for measurement-matched Kermut and ProteinNPT comparisons, with 95% bootstrap confidence bands; the dashed line gives native five-fold Kermut performance. ProteinNPT comparisons include 44 assays and five seeds; Kermut includes 200, 199 and 199 assays at the three budgets. **i**, Mean paired Gaussian-process minus ridge performance on identical ESM2-650M embeddings and measured variants, for ranking and discovery; error bars show 95% assay-bootstrap confidence intervals.

Twenty assay measurements changed the comparison. For each score, we used these measurements to fit an assay-specific linear calibration evaluated on held-out variants. Calibrated ProSST-2048 achieved the highest top-1% recall, 0.204 across 1,065 assay-splits from 213 assays (Supplementary Table 8). Across 20 scores, zero-shot correlation explained less than one-fifth of the variation in calibrated recall (Pearson *r* = 0.44; Fig. 2e), and method rankings changed with measurement budget (Fig. 2f and Supplementary Fig. 3). A ridge combination of leading scores achieved the highest best-top-100 percentile, 0.994, while measured single-mutant additivity provided a biological comparison. Calibrated ProSST-2048 exceeded additivity in 582 of 1,065 matched splits, or 55% (Fig. 2g), showing that additivity was already competitive for discovery at 20 measurements. Calibration, score combinations and additive predictions remained competitive as the budget increased to 50 and 100 measurements (Supplementary Fig. 3).

Increasing the measurement budget shifted the advantage from calibrated priors towards supervised models. Kermut^55^ rose from 0.489 at 20 measurements to 0.633 at 100, while calibrated ProSST-2048 changed from 0.507 to 0.523, leaving calibration ahead at 20 but Kermut ahead at 50 and 100 (Fig. 2h; Supplementary Fig. 4; Supplementary Table 9). ProteinNPT^56^ confirmed the same shift using identical measured variants and held-out candidates, increasing from 0.356 to 0.530 across the same budgets. At 100 measurements, it exceeded calibrated ProSST-2048 by 0.045 (adjusted *q* = 0.0063; Fig. 2h and Supplementary Fig. 5). The balance between pretrained priors and supervised learning therefore changed with measurement depth.

The experimental objective changed model preference even when representation and measurements were held fixed. Using identical ESM2-650M embeddings and measured variants, a Gaussian process outperformed ridge regression in global ranking by 0.029 at 20 measurements and by 0.066 at 100. For top-variant discovery, however, ridge recovered more top-1% variants among its top 100 predictions at 20 measurements, with the two models performing similarly by 50 measurements (Fig. 2i and Supplementary Fig. 4). Improving the ordering of the full library and improving a short testing list thus favoured different models at the smallest budget. Together, these results show that model choice depends on both measurement depth and experimental objective.

### Measurement depth determines the advantage of model priors over additivity

The budget-dependent comparison is particularly consequential for multi-mutants, whose measured component effects provide a biological null. We decomposed double-mutant phenotypes into an additive expectation and an epistatic residual (Fig. 3a and Supplementary Table 10). We evaluated component-resolved doubles, for which both constituent singles were measured, in broad, high-residual, sign-epistasis and reciprocal-sign subsets, and treated variants carrying three or more substitutions separately. High-residual doubles fell in the within-assay top quartile of absolute deviation from additivity. Sign-epistasis candidates showed a reversal in at least one component effect, whereas reciprocal-sign candidates showed both component effects opposite to the combined double-mutant effect relative to the assay reference. These regimes distinguish mutation order from interaction strength and test whether a predictor’s advantage extends beyond recovering component effects (Supplementary Fig. 6).

**Figure 3.**
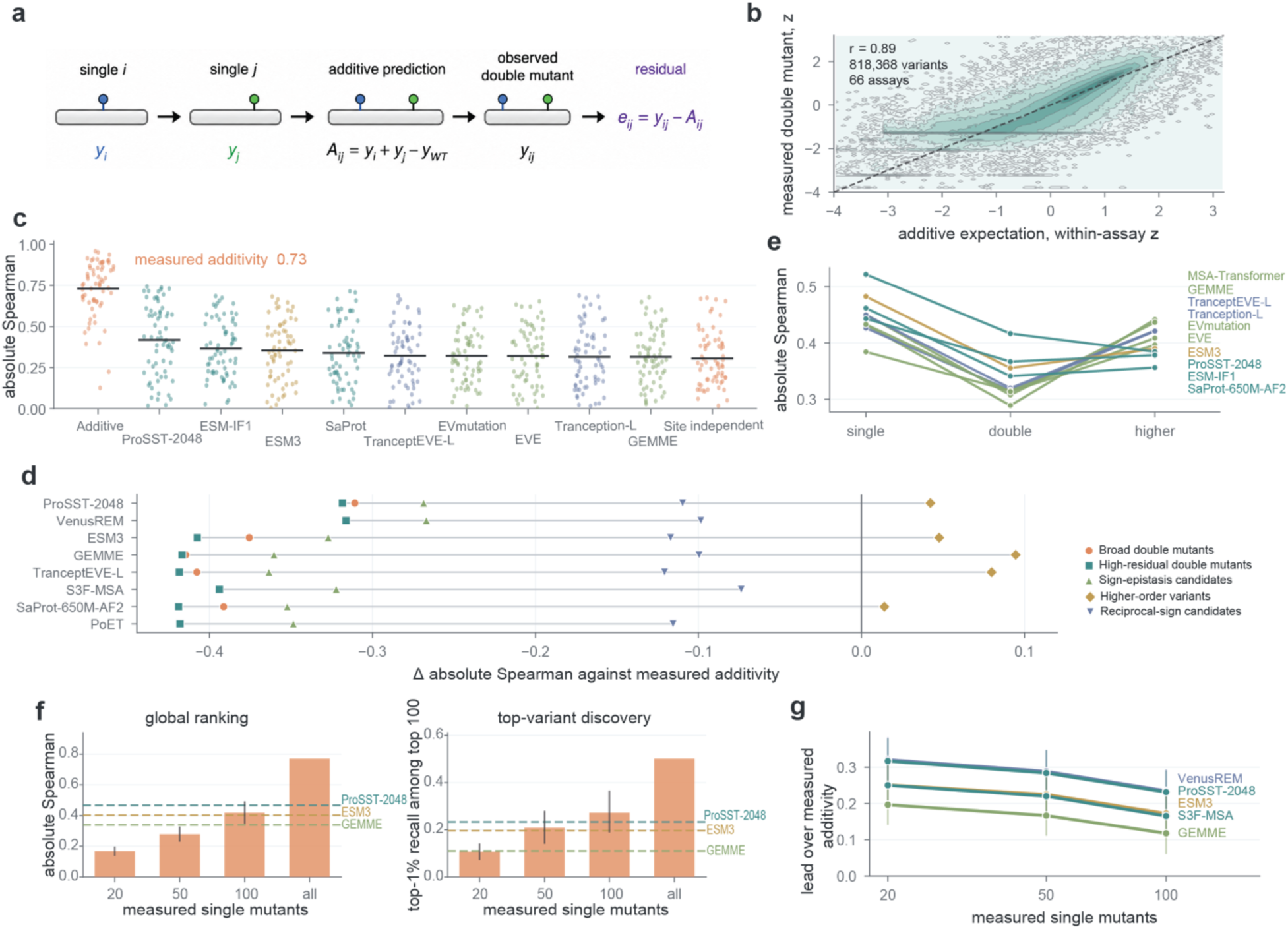
Measurement depth shifts the advantage between measured additivity and model priors. **a**, A double-mutant phenotype decomposed into the additive expectation from its component single mutants and an epistatic residual. The reference phenotype, denoted yWT, is represented by the mean measured single-mutant phenotype in the analysis. **b**, Measured phenotype against additive expectation for 818,368 component-resolved doubles from 66 assays, standardized within assay before pooling; the displayed Pearson correlation summarizes the pooled standardized coordinates. **c**, Absolute Spearman correlation within each assay when all component singles are measured. Points denote assays and horizontal bars denote means; measured additivity reaches 0.730. **d**, Each predictor’s mean gain over additivity by mutation regime. Additivity leads across double-mutant subsets; model means are higher on variants carrying three or more substitutions. **e**, Zero-shot ranking performance across mutation orders, showing the change in method ordering. **f**, Additive performance against component single-mutant measurement depth in a fixed cohort of 31 assays, for ranking and discovery. Bars show assay means and error bars show 95% assay-bootstrap confidence intervals; the all-singles endpoint uses all available component measurements. Dashed lines mark fixed zero-shot references. **g**, Calibrated-prior gains over budget-matched additivity on identical held-out doubles from 65 assays and five seeds, with 95% assay-bootstrap confidence intervals.

With all component single mutants measured, additive predictions closely tracked double-mutant phenotypes, with a pooled Pearson correlation of 0.89 across 818,368 variants from 66 assays after within-assay standardization (Fig. 3b). Across these assays, measured additivity reached a mean absolute Spearman correlation of 0.730, compared with 0.419 for ProSST-2048, 0.355 for ESM3 and 0.316 for GEMME^57^ (Fig. 3c and Supplementary Fig. 6). The advantage persisted among high-residual doubles, where additivity reached 0.734 against 0.417 for the strongest zero-shot predictor (Fig. 3d and Supplementary Table 11). Accurate double-mutant ranking can therefore arise from recovering measured component effects without capturing epistasis. Even within interaction-enriched subsets, claims of interaction learning must exceed a measured additive control.

Mutation order changed the comparison. Measured additivity led across double-mutant subsets, including reciprocal-sign candidates. For variants carrying three or more substitutions, however, the MSA Transformer^58^ ensemble and GEMME reached mean absolute Spearman correlations of 0.442 and 0.437, compared with 0.342 for additivity (Supplementary Fig. 7 and Supplementary Table 12). Zero-shot performance also shifted with mutation order. Every evaluated score lost ranking power on double mutants, and performance on singles and higher-order variants was negatively rank-correlated across the ten leading scores (*r*_s_ = −0.46). The strongest single-mutant predictor ranked eighth on higher-order variants (Fig. 3e). The advantage of measured additivity therefore depends on mutation order and the component information already available. We next varied that information directly to determine how measurement depth changes the comparison.

We next varied the number of measured component singles while holding the assay cohort fixed, isolating measurement depth from differences in assay composition. Across 31 assays with at least 100 component singles, mean absolute Spearman correlation for additivity rose from 0.167 at 20 measurements to 0.277 at 50 and 0.419 at 100, exceeding the fixed zero-shot references for GEMME (0.339) and ESM3 (0.403) but not ProSST-2048 (0.467; Fig. 3f). Discovery crossed earlier. Additivity overtook the zero-shot references for top-1% recall at 50–100 measured component singles, whereas overtaking the leading ranking references required about 500 measurements in the deeper cohort (Supplementary Fig. 8). A short testing list can therefore benefit from measured component effects before those measurements support accurate ranking of the full library. The relevant measurement depth depends on the component coverage needed for the intended experimental decision.

At matched campaign budgets, calibrated priors retained a substantial advantage over additivity. Unlike the component-depth sweep, training singles were sampled from the assay at large rather than specifically from components of the doubles being ranked. Across 65 assays and five seeds, all strategies received identical training singles and held-out doubles. Budget-matched additivity reached Spearman correlations of 0.067, 0.103 and 0.158 at 20, 50 and 100 measurements, respectively. Calibrated VenusREM and ProSST-2048 exceeded additivity by 0.321 and 0.317 at 20 measurements and by 0.235 and 0.232 at 100 (adjusted *q* ≤ 2 × 10^−4^ for both predictors at all three budgets; Fig. 3g and Supplementary Fig. 9). These findings identify complementary information regimes in which calibrated priors retain an advantage at small campaign budgets, while measured additivity becomes stronger as relevant component coverage grows. The value of a model prior therefore depends not only on measurement count but also on which component effects have been measured and the experimental objective.

### Double-mutant measurements benefit models that use mutation identity

Having established the value of measured singles, we asked whether additional double-mutant measurements improve prediction. Within each assay, seed and feature set, we compared models trained with and without additional doubles on shared held-out variants. Adding 20 doubles to 20 measured singles increased held-out Spearman correlation by 0.141 for mutation-identity ridge and by 0.139 for additivity combined with mutation-identity residual features (*n* = 268 assay-seed runs, Fig. 4a and Supplementary Table 13). Gains increased with measurement budget and occurred in 94% of 65 assays at the smallest budget (Fig. 4b, c and Supplementary Fig. 10). Assays selected by a high-epistasis screen gained 0.161, compared with 0.150 among those passed over (Fig. 4d). Additional double-mutant measurements therefore improved prediction when downstream models could exploit mutation identity.

**Figure 4.**
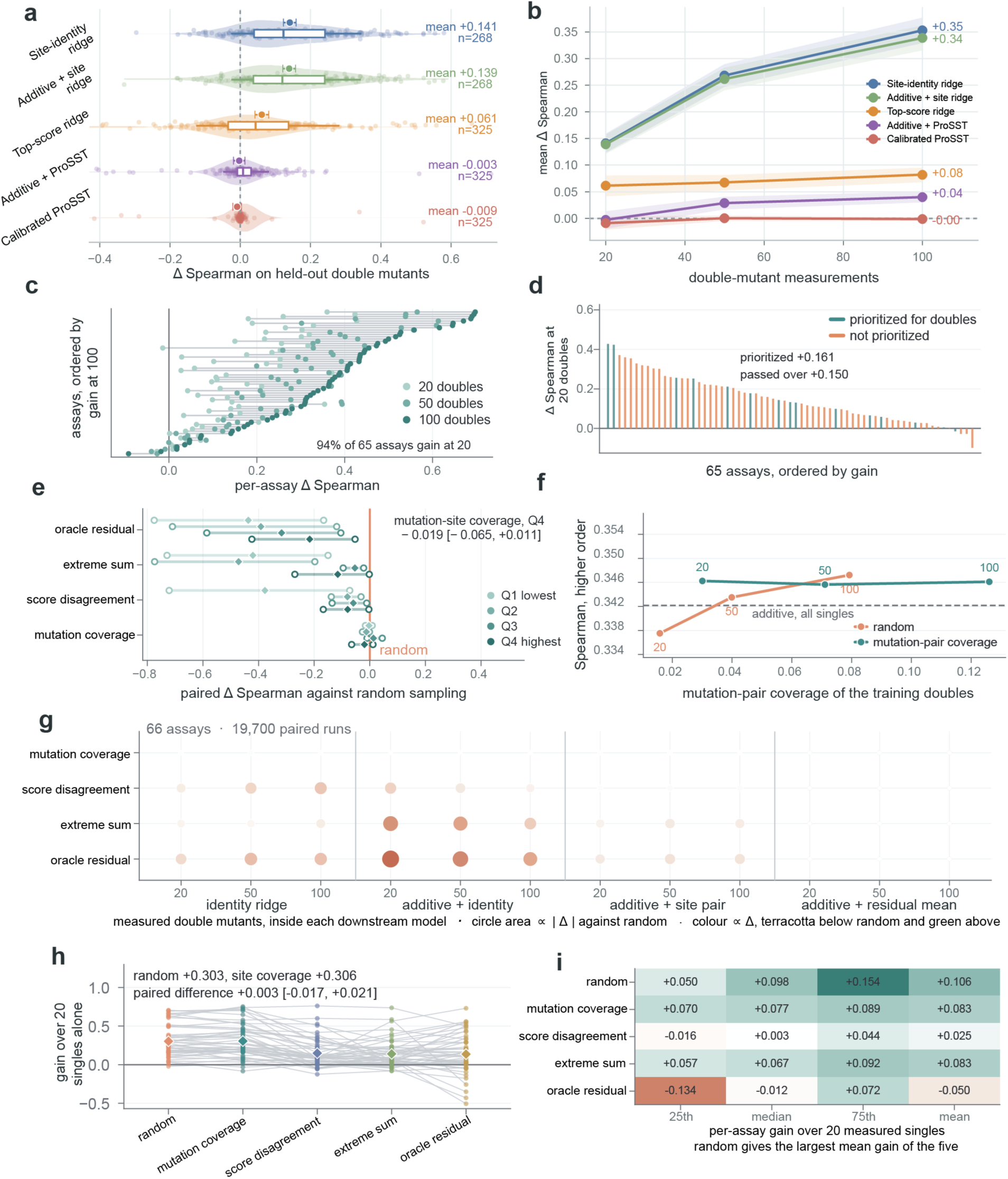
The value of double-mutant measurements depends on the downstream model. **a**, Change in held-out double-mutant Spearman correlation after adding 20 doubles to 20 measured singles. Distributions show assay-seed gains. Boxes span the interquartile range, central lines mark medians and whiskers span the 10th–90th percentiles. Offset circles show means with error bars of ±1.96 standard errors across assay-seed runs; means and sample sizes are labelled. **b**, Mean gain at budgets of 20, 50 and 100 doubles, with bands showing the standard error across evaluable assay-seed runs. **c**, Gains by assay, ordered by the 100-double gain; 94% of 65 assays gain at the smallest budget. **d**, Gains in assays selected or passed over by the high-epistasis rule. **e**, Targeted selection minus random sampling on shared holdouts, grouped by epistasis quartile, with 95% assay-bootstrap confidence intervals. **f**, Higher-order prediction against mutation-pair coverage at each double-mutant budget. **g**, Selection-rule comparisons across four downstream models, three budgets, 66 assays and 19,700 matched runs. Circle area scales with the absolute mean paired difference; terracotta denotes a decrease and green an increase relative to random. Zero has no visible circle. Adding a constant mean residual preserves ranks and therefore yields zero selection-rule differences. **h**, Kermut gains after adding 20 selected doubles; lines connect assay means and diamonds show cross-assay means. **i**, ProteinNPT gains, shown as the 25th percentile, median, 75th percentile and mean. The oracle-residual label denotes selection by measured residual magnitude.

We then asked which doubles to measure. Five selection rules and four downstream models were evaluated on shared holdouts across 66 assays, five seeds and budgets of 20, 50 and 100 doubles. In the highest-epistasis quartile at budget 20, mutation-site coverage yielded a mean paired Spearman difference of −0.019 relative to random sampling for additivity plus mutation identity (Fig. 4e). Selection by extreme additive predictions or model disagreement also often yielded less useful training sets than random sampling.

Greater mutation-pair coverage did not translate into better prediction of higher-order variants. Across 11 assays, mutation-site coverage increased pair coverage 1.6–1.9-fold over random sampling, but its mean absolute Spearman correlation remained near 0.346 across budgets, while random sampling rose from 0.338 at 20 doubles to 0.347 at 100 (Fig. 4f). Across 19,700 matched runs, random sampling attained the highest mean correlation in 10 of 12 model–budget comparisons (Fig. 4g). These results establish random sampling as a demanding reference for assessing whether targeted double-mutant measurements improve downstream prediction.

We extended the selection comparison to Kermut and ProteinNPT, each trained on 20 singles plus 20 selected doubles. Across Kermut assays, random sampling and mutation-site coverage increased Spearman correlation by 0.303 and 0.306, respectively, with a paired difference of 0.003 (adjusted *q* = 0.78; Fig. 4h and Supplementary Fig. 5). Across ProteinNPT assays, random sampling gave the largest mean gain, 0.106, compared with −0.050 to 0.083 for targeted rules (Fig. 4i and Supplementary Fig. 5). These results establish that the value of a double-mutant selection rule must be assessed through its downstream model against random sampling on a shared holdout (Supplementary Table 13).

### Prediction-guided acquisition improves experimental discovery

We extended the measurement-allocation test from fixed training sets to successive acquisition rounds (Supplementary Table 14). We first asked what candidate-level diagnostic scores identify. Across 66 assays, six predictors calibrated on component singles supplied seven measures of prediction level or disagreement (Fig. 5a). Among the ten doubles with the greatest prediction spread, positive-tail epistatic residuals were enriched 4.31-fold (adjusted *q* = 4.00 × 10^−5^), while additive-versus-ensemble differences enriched negative-tail residuals 2.94-fold (adjusted *q* = 4.00 × 10^−5^; Fig. 5b, c and Supplementary Table 15). Prediction spread also enriched the largest absolute residuals 2.90-fold, with enrichment concentrated near the top of each ranking (Fig. 5c). Different disagreement measures therefore identified different forms of model mismatch, but their value for acquisition remained to be established through downstream discovery at matched measurement costs.

**Figure 5.**
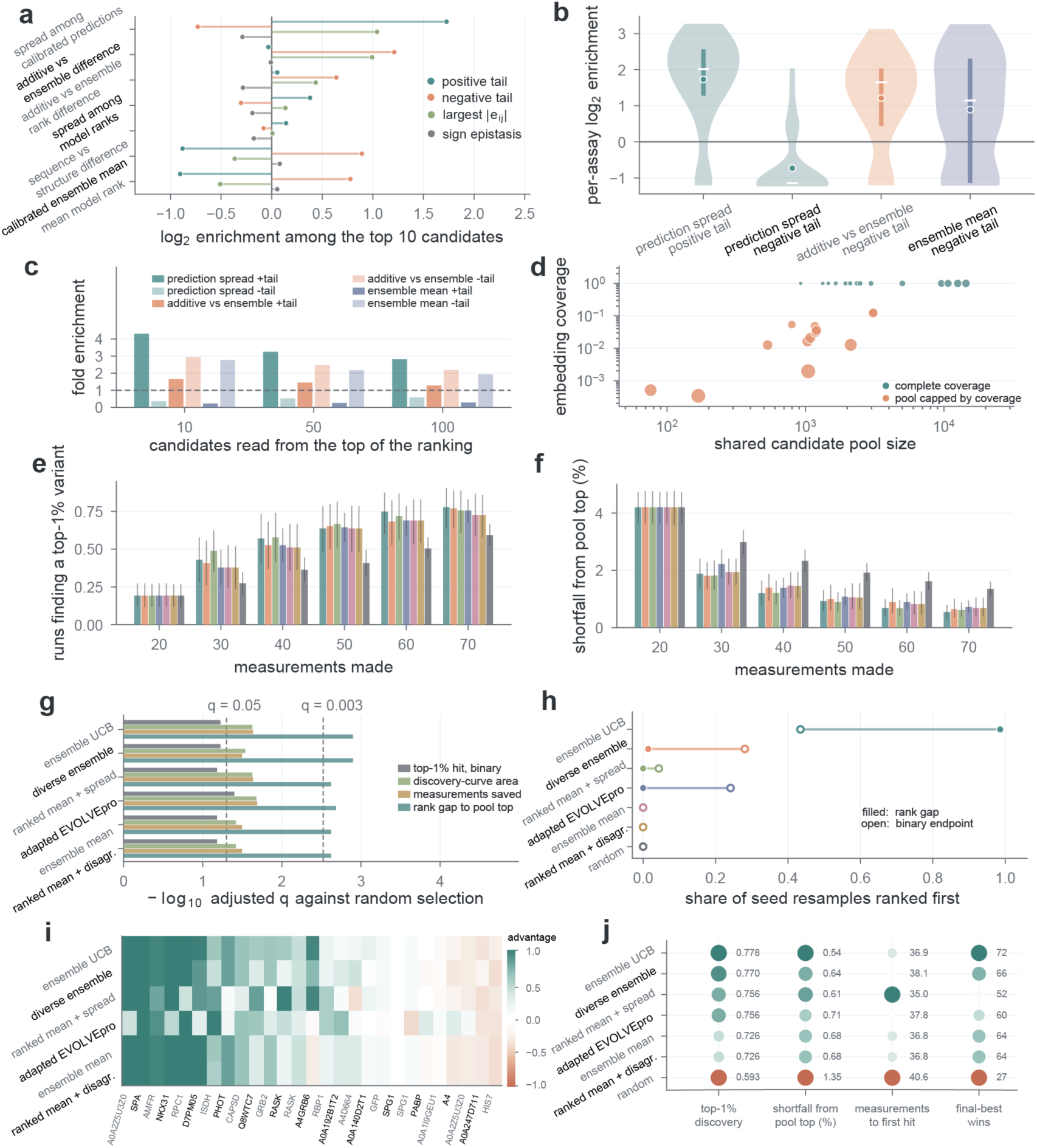
Prediction-guided acquisition improves discovery on a shared candidate pool. **a**, Mean per-assay log2 enrichment of four epistatic-residual classes among the ten highest-ranked doubles for each diagnostic score, across 66 assays. **b**, Distributions of per-assay log2 enrichment for selected contrasts. Circles mark means, white lines medians and thick vertical lines interquartile ranges. **c**, Arithmetic mean fold enrichment at each candidate-list length. **d**, Candidate-pool size against embedding coverage in 27 assays. **e**, Cumulative top-1% discovery from 20 shared initial measurements to 70. **f**, Shortfall of the best measured variant from the pool maximum. In e,f, bars show assay-level means over five seeds and error bars show 95% assay-bootstrap confidence intervals (27 assays; 135 runs per policy). **g**, Prediction-guided policies versus random selection on four endpoints, plotted as −log10 Benjamini–Hochberg adjusted q from two-sided paired sign-permutation tests, adjusted across nine policy-versus-random contrasts separately for each endpoint; dashed lines mark q = 0.05 and 0.003. **h**, Frequency of ranking first under seed resampling, for rank gap and binary discovery. **i**, Per-assay advantage over random selection. **j**, Final outcomes for discovery rate, shortfall, mean measurements to first hit among successful runs and counts of final-best wins. Tied policies each receive a win. Circle areas scale with performance within each metric; labels give the measured values.

Acquisition policies were compared on shared candidate pools from 27 assays, comprising 135 assay-seed runs. Each started from the same 20 measured variants and acquired five batches of ten, with measured phenotypes revealed and policies updated after each batch (Fig. 5d). At the final budget of 70 unique measurements, six prediction-guided policies achieved top-1% discovery rates of 0.726–0.778, compared with 0.593 for random selection. Ensemble upper confidence bound reached 0.778, followed by diversity-aware selection at 0.770 (Fig. 5e, Supplementary Figs. 11 and 12 and Supplementary Table 14). Discovery increased across rounds, and the best measured variants approached the candidate-pool maximum (Fig. 5f). Shared starting variants and candidate pools allowed direct comparison of discovery at the same measurement cost.

All six prediction-guided policies improved final rank gap over random selection (adjusted *q* < 0.003), as well as discovery-curve area and measurements saved (adjusted *q* < 0.05; Fig. 5g). Ensemble upper confidence bound ranked first in 99% of seed resamples on the rank-gap endpoint (Fig. 5h). Its advantage over random held in 19 of 27 assays, and guided policies improved all final decision metrics (Fig. 5i,j). These results show that prediction-guided acquisition can improve experimental discovery, but the value of an acquisition signal must be established through downstream outcomes at matched measurement costs rather than diagnostic enrichment alone.

### An evidence hierarchy yields assay-level recommendations

The preceding comparisons establish an evidence hierarchy for selecting assay-level starting strategies as measurements accumulate. We integrated this evidence into a fixed-priority map using four sequential checks of 20-measurement calibration gain, normalized epistasis, structure-informed gain and alignment depth (Fig. 6a). The first satisfied condition determined the starting strategy, with additive or low-data supervised models as the fallback. Applied to 217 assays, the map grouped assays by the evidence supporting their assignments (Fig. 6b).

**Figure 6.**
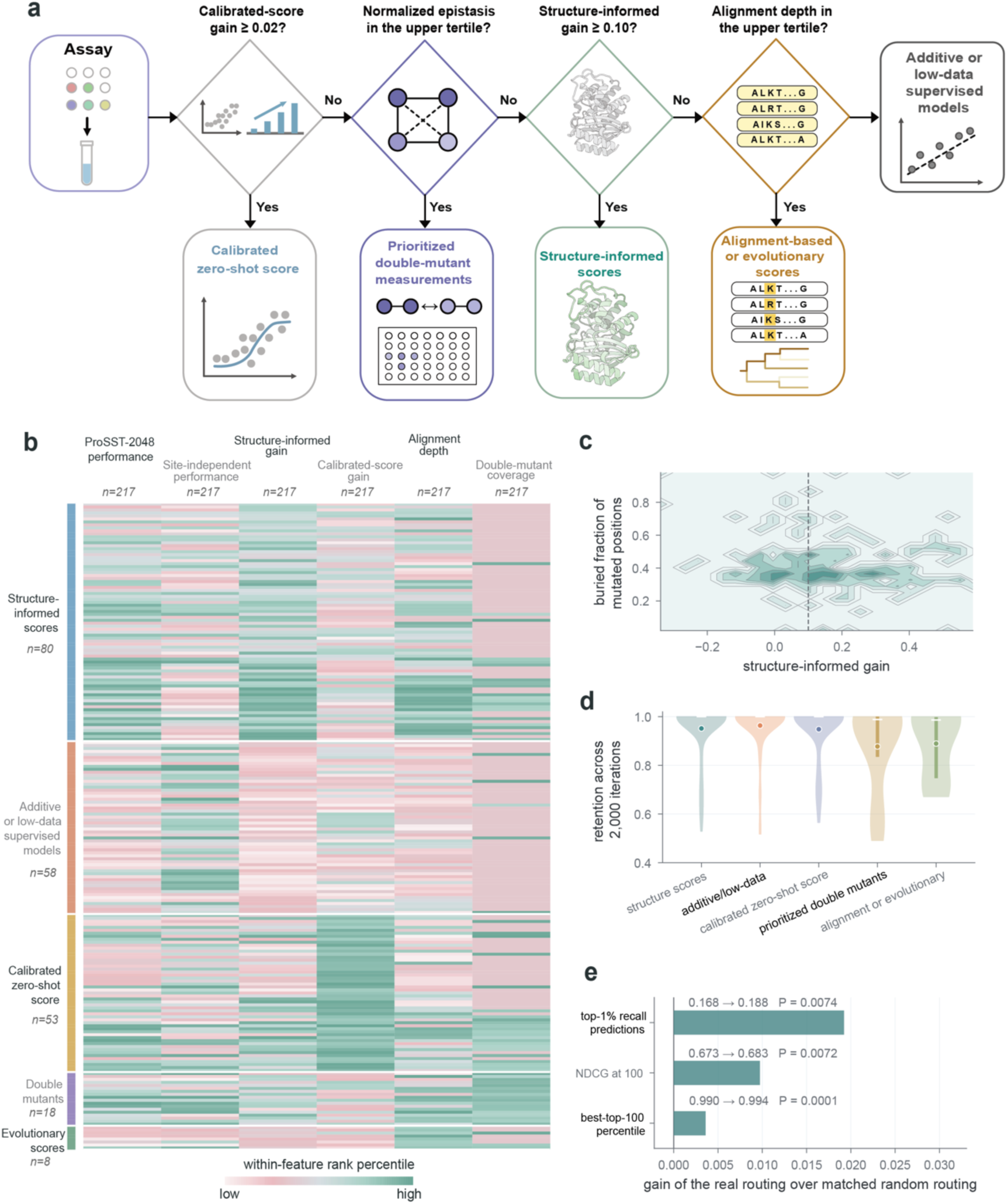
An evidence hierarchy assigns assay-level starting strategies. **a**, Four sequential evidence checks with fixed priority; the first satisfied condition determines the recommendation, with additive or low-data supervised models as the fallback. **b**, Evidence profiles for 217 assays grouped by recommendation. Colours indicate within-feature rank percentiles; group sizes are labelled. **c**, Buried fraction of mutated positions against structure-informed gain. The broad gain range supports routing by measured structural benefit. **d**, Assignment retention across 2,000 protein-clustered perturbation iterations, by recommendation. Circles mark means, white lines medians and thick vertical lines interquartile ranges of per-assay retention. **e**, Real routing versus random routing with the same category sizes across 198 assays with evaluable scoring strategies, for three decision metrics. Bars show mean gains; labels give outcome means and one-sided permutation P values from 10,000 permutations. Recall and normalized discounted cumulative gain evaluate discovery and prioritization; best-top-100 percentile also supplies the calibration-gain rule.

The map assigned 80 assays to structure-informed scores, 58 to additive or low-data supervised models, 53 to calibrated zero-shot scores, 18 to double-mutant measurement and 8 to alignment-based or evolutionary scores (Fig. 6b; Supplementary Tables 17 and 18). Calibration gain most sharply distinguished the groups (Supplementary Fig. 13). Structure-informed gain ranged from −0.36 to 0.61 while the buried fraction of mutated positions remained concentrated in a partially buried band, supporting the use of measured structural benefit for routing (Fig. 6c; Supplementary Fig. 13; Supplementary Table 16). Leave-one-protein threshold re-estimation retained 216 of 217 assignments (99.54%). Across 2,000 protein-clustered perturbations, mean retention ranged from 0.877 for double-mutant measurement to 0.963 for additive or low-data models (Fig. 6d and Supplementary Table 18).

Routing linked assay-level evidence to experimental outcomes. Across 198 assays with evaluable scoring strategies, evidence-based routing increased top-1% recall from 0.168 to 0.188, normalized discounted cumulative gain at 100 from 0.673 to 0.683 and best-top-100 percentile from 0.990 to 0.994 relative to random routing with identical category sizes (all *P* < 0.008; Fig. 6e and Supplementary Table 19). Preserving category sizes tests whether assigning each strategy to a particular assay adds value beyond how frequently that strategy is used. These results show that assay-specific evidence can improve strategy selection and candidate prioritization, connecting model evaluation to experimental design.

## Discussion

Model advantage changes as a campaign acquires information. At matched budgets of 20–100 measurements, leading calibrated priors exceeded budget-matched additivity by 0.23–0.32 Spearman correlation units. As relevant component singles accumulated within fixed assay cohorts, measured additivity overtook zero-shot references at 50–100 measurements for top-variant discovery, whereas surpassing the leading global-ranking references required about 500 in the deeper cohort. These comparisons identify complementary information regimes, with the crossover depending on component coverage and experimental objective rather than measurement count alone. The findings connect model selection to experimental design by showing that the value of a model prior depends on both the biological information already measured and the decision it must support.

Measured component effects also change the standard for evaluating model predictions. With component singles available, additivity reached a mean absolute Spearman correlation of 0.730 for double-mutant ranking, compared with 0.419 for the strongest zero-shot predictor. Its advantage persisted within interaction-enriched subsets, showing that accurate double-mutant ranking can arise from recovering component effects without capturing epistasis. Claims of interaction learning must therefore be evaluated against an explicit additive control with a stated amount of component information. The same principle extends to model adaptation, where zero-shot correlation explained less than one-fifth of the variation in discovery after assay-specific calibration. A model’s contribution must be assessed relative to the biological information already measured and the experimental decision it supports.

For experimental planning, measurement count alone does not determine the information available to a predictor. The same number of measured singles can provide different coverage of the component mutations in the candidate pool, changing the relative value of additivity and model priors. A short testing list can benefit from measured component effects before those measurements support accurate ranking of the full library. Predictor choice should therefore be revisited as component coverage grows, using the intended experimental objective to determine whether newly acquired measurements change the preferred strategy.

Additional measurements extend the comparison from predictor choice to experimental allocation. Random sampling provided a strong reference for selecting double-mutant training data to improve prediction, whereas prediction-guided acquisition improved top-variant discovery across successive batches. These findings reflect different experimental objectives rather than conflicting sampling preferences. The value of a measurement must therefore be established through the downstream model and decision it supports, with selection strategies evaluated against appropriate controls at the same experimental cost. This distinction connects the assessment of predictive accuracy to the allocation of measurements in an engineering campaign.

The assay-level evidence map translates these principles into starting strategies. Across assays with evaluable scoring strategies, evidence-based routing improved discovery and prioritization relative to random routing with identical category sizes. This shows that assay-specific evidence can inform which strategy to use, beyond comparisons of predictor accuracy alone. Our analyses are retrospective and limited to substitution landscapes. Extending this approach to insertions, deletions and de novo design will require appropriate biological controls and explicit measurement costs. By establishing when a predictor adds value, when measured information changes that advantage and which further measurements improve the intended outcome, decision benchmarking connects model evaluation to experimental design.

## Methods

### Benchmark design and data

We constructed decision benchmarking from ProteinGym substitution assays. Each method was evaluated as an assay-level strategy operating under a fixed measurement budget. A strategy received an assay-specific candidate pool, optional precomputed zero-shot scores, a measured training set and reference controls that used the same available information. It returned a ranked list, calibrated predictor, fitted supervised model, double-mutant measurement panel, acquisition batch or assay-level recommendation. We used substitution assays because they provide measured candidate pools, component mutations and precomputed variant scores needed for matched retrospective comparisons. Budgets of 20, 50 and 100 measurements represent early experimental stages in which model choice can materially change which variants are tested. Performance comparisons were computed within assay or assay-split replicate and then averaged across eligible units, giving each assay equal weight within the stated comparison. Figure 3b separately displays pooled coordinates standardized within assay; its Pearson correlation describes that display. Within-assay metrics underpin the model-performance contrasts in the main figures. Supplementary Table 11 also reports pooled reciprocal-sign correlations. Full methodological specifications are provided in Supplementary Methods.

We analysed 217 ProteinGym substitution assays containing 2,465,767 measured variants: 696,311 single mutants, 826,245 double mutants and 943,211 variants with three or more substitutions. Continuous ProteinGym scores were used as the primary phenotype. Binary labels were used only for assays with available ProteinGym binary annotations. Variant strings were parsed into component substitutions, mutation order and mutation sites using a single rule throughout the study. Official ProteinGym metadata were used for assay-level annotation, multiple-sequence-alignment depth and structural-context analyses, but not as training features for the main models.

### Zero-shot predictors and design metrics

We evaluated 95 precomputed zero-shot predictors as prior scores. These scores included sequence language models, MSA-based models, evolutionary models, structure-informed models, hybrid or retrieval-augmented models and simple variant-effect controls. Method families were used only for interpretation; each score column was evaluated independently. Variants with missing or invalid scores were excluded only from the corresponding method– assay comparison.

Zero-shot performance was summarized by within-assay Spearman correlation between predictor scores and measured phenotypes. Spearman correlation measures how well a method orders the full candidate set; absolute Spearman was used for orientation-invariant summaries. Design-oriented metrics used the supplied higher-is-better score direction without per-assay sign correction. Top-1% recall among the top 100 predictions measures how many of the assay’s highest-scoring 1% of variants are recovered in a 100-variant testing list. Top-20 enrichment measures the fold increase over random selection in recovering those top-1% variants among the 20 highest-ranked candidates. Best-top-100 percentile records the percentile of the best measured variant among the top 100 predictions and therefore asks whether a short predicted list contains at least one very strong candidate. Normalized discounted cumulative gain rewards placing high-scoring measured variants near the top of the list. Where ProteinGym binary labels were available, area under the receiver-operating-characteristic curve measured binary discrimination and Matthews correlation coefficient summarized thresholded classification. These metrics represent different experimental objectives and were not combined into one universal score.

### Low-measurement calibration and supervised baselines

Few-measurement experiments used deterministic training splits of 20, 50 and 100 variants per assay, with five split replicates. We considered three training-set designs: single-mutant-only, mixed-variant and single-plus-double splits. Training and held-out variants were disjoint within each assay-split; each measured variant appeared once in its training set. General low-measurement evaluations used up to 20,000 held-out candidates per assay-split, sampled deterministically without phenotype information; the external-model comparisons used their specified common candidate pools.

Each calibrated zero-shot model fitted an assay-specific intercept and slope relating one precomputed score to training-set phenotypes. A single-mutant additive model estimated substitution effects from measured single mutants and predicted multi-mutant variants by summing observed component effects, with unseen component substitutions contributing zero relative to the baseline. Additive-plus-score models combined this additive prediction with a calibrated zero-shot score. Score-combination models fitted ridge regression to a fixed panel of nine leading zero-shot scores: VenusREM, ProSST-2048, S3F-MSA, S2F-MSA, PoET, RSALOR, ESM3, GEMME and SaProt-650M-AF2. A second combination model used mutation order, the summed measured single-mutant effect and eight zero-shot scores, VenusREM, ProSST-2048, S3F-MSA, S2F-MSA, PoET, ESM3, GEMME and TranceptEVE-L, as features. Both panels were fixed before any budgeted comparison.

We evaluated ridge regression, lasso, elastic net, random forest, histogram gradient boosting, XGBoost and a Gaussian-process surrogate. Hyperparameters were fixed before comparison. Models were fitted separately for each assay, seed, budget and training-set design. Official Kermut^55^ and ProteinNPT^56^ outputs were summarized in the ProteinGym random, modulo and contiguous five-fold settings. Kermut was also fitted at budgets of 20, 50 and 100 using its composite mutation-kernel Gaussian process, ESM2-650M representations, the ESM2-650M zero-shot score, ProteinMPNN conditional probabilities and structural coordinates. Ridge regression, histogram gradient boosting, XGBoost and a Gaussian-process surrogate were fitted to the same ESM2-650M mean-pooled embeddings and measured variants as capacity controls.

ProteinNPT was retrained on 44 assays with complete runs for five deterministic seeds and budgets of 20, 50 and 100 measurements. Proteins were no longer than 400 residues. Candidate pools retained every training variant and were capped at 2,000 variants by phenotype-independent sampling. ProteinNPT and all comparison models were evaluated on identical training and held-out candidate identities. Additive-control Spearman correlations were reported for splits with variable predictions; mixed-variant splits with constant additive predictions were excluded from that metric.

### Additive controls and multi-mutant regimes

For double-mutant analyses, the biological reference was a measured single-mutant additive control. A double mutant was component resolved if both corresponding single mutants were measured in the same assay. The mean phenotype of measured single mutants served as the assay reference, providing a consistent proxy across assays without uniformly available wild-type measurements. The additive expectation was the sum of the two-component single-mutant phenotypes minus this assay reference. With both component effects measured, changing the reference translates every additive expectation by the same constant and preserves their rank correlation. The epistatic residual was the observed double-mutant phenotype minus the additive expectation. A positive residual means the measured double mutant performed above the additive expectation, whereas a negative residual means it performed below that expectation. Normalized epistasis summarized an assay by its median absolute epistatic residual divided by the interquartile range of the observed double-mutant phenotypes in the same assay, so that the measure is comparable across assay scales.

Strict residual analyses used the 66 assays containing at least 100 component-resolved double mutants, 818,951 variants in total. Figure 3b displays the 818,368 variants within the plotted range after within-assay standardization; its Pearson correlation was calculated on those displayed coordinates. Broad double mutants included all eligible component-resolved variants. High-residual double mutants were the within-assay top quartile by absolute epistatic residual. A sign-epistasis candidate was a double mutant in which the apparent direction of at least one component mutation changed when the second mutation was present. Reciprocal-sign candidates required both component single mutants to have a direction opposite to the combined double mutant relative to the assay reference. Higher-order variants contained three or more substitutions. Each regime was evaluated separately. Gain over additive was the difference in absolute Spearman correlation between the predictor and additive control within the same assay and regime.

We varied the component information available to the additive control by subsampling measured single mutants at each depth while holding the assay cohort fixed, so that measurement depth was not confounded with assay composition. Cohorts were defined by requiring assays to contain at least the target number of component single mutants. Within each cohort, a specified number of measured single mutants was drawn without replacement at each depth, the additive expectation was rebuilt under the rule above with unobserved components contributing zero relative to the baseline, and both the ranking metric and the top-variant discovery metric were recomputed on every component-resolved double mutant in the assay. At each depth, the baseline and observed component effects were estimated from the sampled singles. Cohorts contained 31 assays with at least 100 component singles and 11 assays with at least 500, and five draws were averaged within assay before cohort means were calculated.

A budget-matched comparison used measured single mutants for training and shared held-out double mutants for evaluation. Across 65 assays, five deterministic seeds and budgets of 20, 50 and 100 measured single mutants, the additive control, each calibrated zero-shot score and their combination received identical training variants and identical held-out candidates. The priors were the eight-score panel above; Benjamini–Hochberg correction covered the eight predictors and three contrasts separately at each budget.

For single-to-multi generalization, Kermut was trained on 20, 50 or 100 measured single mutants and ProteinNPT on 20. Both were evaluated separately on held-out double mutants and variants with three or more substitutions. Kermut used candidates with all required model inputs. ProteinNPT used phenotype-independent candidate pools capped at 800 variants. Spearman correlation was computed within each assay and seed whenever held-out predictions and measurements were variable.

### Double-mutant measurement and active learning

We first compared models trained with and without measured double mutants within the same assay, seed, budget and feature set. We then compared double-mutant selection rules using a common held-out set shared across every policy and budget within an assay and seed. The held-out set contained 20% of eligible double mutants, with a minimum of 50 and a maximum of 10,000, while retaining enough acquisition candidates for the largest feasible budget. All available component single mutants were used as side information for the additive control and assay-level ProSST-2048 calibration and were not counted toward the double-mutant measurement budget. Budgets of 20, 50 and 100 double mutants were tested with five seeds. Selection rules were random sampling, mutation-site coverage, extreme additive prediction, disagreement between additive and calibrated ProSST-2048 predictions and a retrospective largest-residual diagnostic. Four downstream models were used: an additive prediction shifted by the mean measured residual, an additive prediction with residuals estimated for observed site pairs, ridge regression on mutation-identity indicators, and the additive prediction combined with mutation-identity ridge residuals. Mutation-site coverage prioritized previously unrepresented substitutions, using newly covered sites to break ties. Extreme additive selection ranked candidates by absolute deviation from the acquisition-pool median additive prediction. The largest-residual rule ranked acquisition candidates by measured residual magnitude and served as a retrospective diagnostic; the shared holdout was reserved for evaluation. For higher-order transfer, a separate coverage rule ranked measured doubles by the frequency of their substitution pairs in the target higher-order sequences.

The external selected-double comparison used the same five selection rules with recent supervised models. Kermut was evaluated across 51 assays with complete embeddings, zero-shot scores, structural coordinates and conditional probabilities. ProteinNPT was evaluated on four sampled assays with complete model resources. Every run started from the same 20 single mutants, added 20 selected double mutants and used a shared held-out double-mutant set within each assay and seed. Gains were computed against the corresponding model trained on the same 20 single mutants alone.

Diagnostic scores were evaluated before acquisition. Six zero-shot predictors, ProSST-2048, ESM3, S2F-MSA, S3F-MSA, PoET and VenusREM, were linearly calibrated to each assay using unique component single mutants. Seven candidate-level summaries were then computed: the calibrated ensemble mean and standard deviation, the difference between the additive prediction and the ensemble mean, sequence-versus-structure and additive-versus-ensemble rank disagreement, and the mean and standard deviation of calibrated ranks. Within each assay, double mutants were ranked by each summary and the top 10, 50 and 100 candidates were taken. Four events were tested: the upper and lower tenths of signed residuals, the largest tenth of absolute residuals and sign-epistasis candidates. Enrichment was the event frequency in the selected set divided by its frequency among all eligible double mutants in the same assay. Means and 95% confidence intervals came from assay-level bootstrap resampling. Inference tested smoothed log2 enrichment, adding 0.5 to event counts and 1 to totals in both selected sets and assay backgrounds before taking the ratio. Benjamini–Hochberg correction covered all score, set-size and event combinations.

Retrospective active learning used a shared candidate pool for each of 28 representative assays. Candidate pools intersected measured ProteinGym variants with available ESM2-650M embeddings and were capped by deterministic phenotype-independent sampling when necessary. One assay had fewer than 70 eligible variants, leaving 27 assays and 135 assay-seed runs. Every policy started from the same 20 variants and selected five batches of ten. After each batch, the measured ProteinGym phenotypes were revealed and the policy was updated. Policies included random selection, calibrated ensemble-mean ranking, ensemble upper confidence bound, diversity-aware selection, rank combinations of predicted score with between-model spread or additive disagreement, and the EVOLVEpro random-forest top-ranking rule^46^. Adapted implementations of further published acquisition rules^59^ were evaluated on the same shared pools and are specified in Supplementary Methods. EVOLVEpro used ESM2-650M embeddings. ALDE, the active learning-assisted directed evolution rule^59^, used binary mutation-identity features and a gradient-boosted ensemble. The primary outcome was the fraction of runs measuring at least one true top-1% variant in the shared candidate pool. Three further endpoints were recorded for every run. Discovery-curve area was the mean of that indicator across the five acquisition batches. Measurement efficiency was defined as the negative restricted mean number of measurements to the first true top-1% variant, truncated at 70, so that a positive policy-minus-random difference directly represents measurements saved relative to random selection. Rank gap to the top of the pool was the negative base-10 logarithm of one plus the number of pool variants ranked above the best variant a run had measured after the final batch. All four endpoints were oriented so that higher values indicated better performance. Descriptive first-hit summaries report measurement counts among successful runs: Figure 5j shows means and Supplementary Figure 12c shows individual runs and medians. Final-best wins count each policy attaining the best measured phenotype within an assay and seed, including ties.

### Assay-level recommendation map and statistics

We converted measured assay evidence into a fixed-priority assay-level recommendation map. Inputs combined held-out performance after calibration, component-resolved epistasis, assay-level zero-shot performance and alignment depth. Assays were assigned to calibrated zero-shot scoring if the best 20-measurement calibrated score improved best-top-100 percentile over the additive baseline by at least 0.02, the calibrated-score gain. Otherwise, assays whose normalized epistasis reached the empirical two-thirds quantile with lower interpolation, realized at 0.6585, and that contained measured double mutants, were assigned to prioritized double-mutant measurement. This recommendation allocates measurements to double mutants; selection policies are evaluated separately against random sampling. Remaining assays were assigned to structure-informed scoring if ProSST-2048 exceeded the site-independent score by at least 0.10 absolute Spearman correlation, the structure-informed gain, to alignment-based or evolutionary scoring if effective alignment depth divided by protein length reached the upper tertile of the assay distribution, realized at 95.02, and to additive or low-data supervised models otherwise. If multiple conditions applied, the earliest satisfied rule determined the recommendation.

Statistical summaries are provided in Supplementary Table 19. Cross-assay estimates were reported as mean over assay-level or assay-split-level values, depending on the analysis. Seeds were averaged within assay before bootstrap confidence intervals and paired tests for the selection and active-learning comparisons. The paired inferential analyses used 5,000 deterministic bootstrap resamples. Display-specific resampling counts are stated in the figure legends. Selected paired comparisons used two-sided sign-permutation tests with Benjamini– Hochberg correction. Figure 3e uses 2,000 assay-bootstrap resamples for its plotted intervals; Figure 4a shows mean ±1.96 standard errors and Figure 4b shows mean ±1 standard error across assay-seed runs. Decision-map stability was assessed in two ways. First, thresholds derived from assay distributions were recomputed after leaving out one protein at a time. Second, 2,000 protein-clustered iterations jointly resampled proteins, recomputed the epistasis and alignment-depth thresholds and varied the fixed calibration-gain and structure-gain thresholds by ±20%. The association between zero-shot ranking and post-calibration utility was a Pearson correlation between mean zero-shot absolute Spearman correlation and mean 20-measurement top-1% recall in the mixed-variant design, across method-level means, one point per zero-shot score, for the 20 scores evaluated at that budget. For the four scoring categories, each category was represented by the method with the highest cross-assay mean for the evaluated endpoint within its designated family, with additivity representing the additive category. Of 217 assays, 18 received a double-mutant-measurement recommendation, an experimental action. The remaining 199 were eligible for scoring-strategy comparison; complete outcomes were available for 198 assays. Best-top-100 percentile also contributed to the calibration-gain rule, while recall and normalized discounted cumulative gain supplied distinct outcome measures. Routing was compared against a size-preserving random control in which the recommendation labels were permuted across assays while the number of assays in each recommendation category was held fixed, and the realized outcome was recomputed for 10,000 such permutations. One-sided permutation tests compared the observed routing outcome with this random-routing distribution. For the four active-learning endpoints in Figure 5g, Benjamini–Hochberg correction was applied separately within each endpoint to nine policy-versus-random contrasts. The two-outcome acquisition comparisons in Supplementary Figures 11d and 12d were adjusted jointly across 36 contrasts; the uncertainty-versus-mean comparisons were adjusted across eight contrasts. Benjamini–Hochberg adjusted values are reported as adjusted *q*. Missing values were not imputed. All analyses were retrospective computations on existing ProteinGym measurements and precomputed scores.

## Author contributions

W.D. conceived the study and developed the methodology. W.D., C.L., Y.L., H.G. and Z.W. all contributed to the conception of the work. W.D. and Y.L. performed the analyses. W.D. interpreted the results and wrote the initial manuscript. C.L., H.G. and Z.W. revised the manuscript.

